# Xylazine co-self-administration suppresses fentanyl consumption during self-administration and induces a unique sex-specific withdrawal syndrome that is not altered by naloxone in rats

**DOI:** 10.1101/2023.05.17.541158

**Authors:** Shailesh N. Khatri, Safiyah Sadek, Percell T. Kendrick, Emma O. Bondy, Mei Hong, Sally Pauss, Dan Luo, Thomas E. Prisinzano, Kelly E. Dunn, Julie A. Marusich, Joshua S. Beckmann, Terry D. Hinds, Cassandra D. Gipson

**Affiliations:** Department of Pharmacology and Nutritional Sciences, University of Kentucky, Lexington KY; Center for Pharmaceutical Research and Innovation, College of Pharmacy, University of Kentucky, Lexington, KY; Psychiatry and Behavioral Sciences Department, Johns Hopkins University, Baltimore, MD; Center for Drug Discovery, RTI International, Research Triangle Park, NC; Department of Psychology, University of Kentucky, Lexington, KY

**Keywords:** Xylazine, fentanyl, withdrawal, self-administration, sex differences

## Abstract

Prescription and illicit opioid use are a public health crisis, with the landscape shifting to fentanyl use. Since fentanyl is 100-fold more potent than morphine, its use is associated with a higher risk of fatal overdose that can be remediated through naloxone (Narcan) administration. However, recent reports indicate that xylazine, an anesthetic, is increasingly detected in accidental fentanyl overdose deaths. Anecdotal reports suggest that xylazine may prolong the fentanyl “high”, alter the onset of fentanyl withdrawal, and increase resistance to naloxone-induced reversal of overdose. To date no preclinical studies have evaluated the impacts of xylazine on fentanyl self-administration (SA; 2.5 μg/kg/infusion) or withdrawal to our knowledge. We established a rat model of xylazine/fentanyl co-SA and withdrawal and evaluated outcomes as a function of biological sex. When administered alone, chronic xylazine (2.5 mg/kg, IP) induced unique sex-specific withdrawal symptomatology whereby females showed delayed onset of signs and a possible enhancement of sensitivity to the motor-suppressing effects of xylazine. Xylazine reduced fentanyl consumption both male and female rats regardless of whether it was experimenter-administered or added to the intravenous fentanyl product (0.05. 0.10, and 0.5 mg/kg/infusion) when compared to fentanyl SA alone. Interestingly, this effect was dose-dependent when self-administered intravenously. Naloxone (0.1 mg/kg, SC) did not increase somatic signs of fentanyl withdrawal, regardless of the inclusion of xylazine in the fentanyl infusion in either sex; however, somatic signs of withdrawal were higher across timepoints in females after xylazine/fentanyl co-SA regardless of naloxone exposure as compared to females following fentanyl SA alone. Together, these results indicate that xylazine/fentanyl co-SA dose-dependently suppressed fentanyl intake in both sexes, and induced a unique withdrawal syndrome in females which was not altered by acute naloxone treatment.

## Introduction

Prescription and illicit opioid use are a recognized public health crisis (Mattson et al., 2021), and heroin is being rapidly replaced with illicitly-manufactured fentanyl (Antoine et al., 2021), a synthetic μ-opioid receptor (MOR) agonist that is 100-fold more potent than morphine (Pasternak & Pan, 2013). Fentanyl is associated with a profound increase in fatal overdose cases (Wilson et al., 2020). Naloxone (Narcan™) is a MOR antagonist used clinically to reverse fentanyl’s occupancy of brain MORs (Kang et al., 2022), reversing opioid overdose and precipitating opioid withdrawal (Kosten & Baxter, 2019; Rzasa Lynn & Galinkin, 2018). Recently, xylazine, an adrenergic α_2a_ receptor (A2aR) agonist sold as a veterinary anesthetic, has been increasingly detected in more fentanyl-positive urine screens and fentanyl overdose deaths (Nunez et al., 2021). Anecdotal reports from persons who inject drugs (PWIDs) suggest that xylazine may prolong the fentanyl “high” and delay the onset of fentanyl withdrawal while also reducing naloxone’s efficacy for overdose reversal (Friedman et al., 2022).

As noted above, PWIDs report that xylazine is increasingly evident in illicitly manufactured fentanyl (Reyes et al., 2012), a combination termed “tranq dope” (Friedman et al., 2022). Xylazine adulteration of fentanyl in persons and/or drug supply has been detected in numerous regions throughout the United States and Canada (Bowles et al., 2021; Friedman et al., 2022; Johnson et al., 2021; Nunez et al., 2021). In 2019, xylazine was detected in 31% of unintentional heroin and/or fentanyl overdose deaths and 25% of drugs seized by the Food and Drug Administration (Johnson et al., 2021). Users report that tranq dope is “sought after,” “extends the high [of fentanyl],” solves the “problem” of the “short legs” of illicit fentanyl (i.e., prolongs the duration of fentanyl effects), and decreases the severity of fentanyl withdrawal symptoms (Friedman et al., 2022). However, the health risks associated with tranq dope are also exceptionally high. They include increased risk of overdose death, inability of naloxone to reverse xylazine effects, and medical consequences such as necrotizing skin and soft tissue damage (Friedman et al., 2022). Thus, it is critical to determine the neurobehavioral mechanisms underlying this risky and prevalent pattern of polysubstance use.

Preclinically, studies to date have focused solely on fentanyl self-administration (SA) without the addition of xylazine (e.g., Bardo et al., 2022; Fragale et al., 2020; Hammerslag et al., 2020; Malone et al., 2021) and the only paper to our knowledge that has examined xylazine intravenously did so in combination with ketamine for the purpose of prolonged anesthesia (Linsenmeier et al., 2020). Thus, we conducted a series of behavioral and physiological experiments to examine the following goals. Experiment 1 determined whether xylazine induced its own unique withdrawal syndrome, and Experiment 2A determined if xylazine pretreatment altered fentanyl SA. We then established a working rodent model of intravenous xylazine and fentanyl SA delivered concurrently in the same infusion that was used to examine whether pretreatment with xylazine changed fentanyl SA (Experiment 2C), which was verified using pharmacokinetic (PK) evaluation of intravenous xylazine dosing (Experiment 2B). We also determined whether xylazine changed the trajectory of fentanyl withdrawal and/or occluded the ability of naloxone to precipitate withdrawal overall and in a sex-specific manner.

## Methods

### Subjects

#### Behavioral Experiments (Experiments 1, 2A, and 2C)

92 Long Evans rats (46 male, 46 female) were purchased from Envigo or utilized as wildtypes from an in-house CX3CR1-cre rat breeding colony (LE-Tg(Cx3cr1-cre/ERT2)3Ottc), bred on a Long Evans background for the behavioral studies (breeders initially purchased from the Rat Resource and Research Center). Male rats were 225-250 g and female rats were 200-225 g upon initiation of experimentation and were individually housed on a 12-hour reverse light cycle with ad libitum access to food and water prior to experimental procedures. Animals were handled daily and no animals were excluded from the studies. All animal use practices for behavioral studies were approved by the Institutional Animal Care and Use Committee (IACUC) of the University of Kentucky (UK; Protocol #2020-3438).

#### PK Study (Experiment 2B)

In vivo PK analysis was conducted at WuXi Apptec Co. Ltd. (Shanghai, China). Ten male Sprague Dawley rats were purchased from Vital River Laboratory Animal Technology Co., Ltd. (Beijing, China). Rats were 6 weeks old (290 ± 10 g) upon arrival and were fasted prior to experimentation. No rats were excluded from the analyses.

#### Study Drugs

Xylazine (injectable solution) was purchased from Covetrus (Portland, ME). Fentanyl HCl was gifted from the National Institute on Drug Abuse (NIDA, Bethesda, MD) through the NIDA Drug Supply Program. Naloxone HCl was purchased from Sigma Aldrich (St. Louis, MO).

Xylazine Injection Protocol (Experiment 2A). In Experiment 2A, rats (3/sex/condition) were administered repeated xylazine (2.5 mg/kg, intraperitoneal [IP]) or saline (body weight, IP) for 8 consecutive days. This dose was chosen because it is subanesthetic and there were no prior studies to inform repeated systemic xylazine dosing. Following the 8^th^ day, rats underwent observation for somatic signs of xylazine withdrawal (described below).

Intravenous SA (Experiments 2A, 2C). Jugular vein catheter surgery was conducted with all rats in the SA experiments. Rats were anesthetized with intramuscular (IM) ketamine (80-100 mg/kg) and xylazine (8 mg/kg), and aseptic surgical techniques were utilized. See (Maher et al., 2022) for additional surgical details. Subgroups of rats then underwent either fentanyl SA with xylazine or saline injections (Experiment 2A), or underwent fentanyl SA or xylazine/fentanyl SA procedures (Experiment 2C).

Naloxone Treatment (Experiment 2C). Rats underwent either naloxone HCl (0.1 mg/kg, SC), a dose that has been shown to inhibit morphine-induced place preference and reverse fentanyl-induced respiratory depression in rats (Karimi et al., 2011; Shaykin, 2022), or vehicle immediately following the 9^th^ session of xylazine/fentanyl SA. Somatic signs of withdrawal were then measured across time points as described below.

### Somatic Signs of Withdrawal (Experiments 1, 2C)

Rats underwent observational testing for somatic signs of withdrawal consistent with our prior publication (Gipson et al., 2020). Briefly, sixteen spontaneous signs of withdrawal were measured, including digging, jumping, rearing, grooming, diarrhea, piloerection, genital licks, teeth chatters, chewing, ptosis, escape attempts, eye blinks, foot licks, writhing, head shakes, and body shakes at 0, 24, 48, 120, 240, and 360 hrs post xylazine injection (Experiment 1) or 0, 45 min, 24 hr, 48 hr, 72 hr, and 168 hr, post fentanyl or xylazine/fentanyl SA (Experiment 2C). Rats were placed in a clear Plexiglas chamber (9.6 × 9.6 × 14.6in; L x W x H) and recorded for 10 minutes after 5 minutes of habituation. Experimenters were blinded to group and timepoint and scored withdrawal signs from the video recordings using a standardized scoring protocol (Gipson et al., 2020).

### Operant Behavior (Experiments 2A, 2C)

#### Operant Conditioning Chambers

28 sound-attenuating chambers with ventilation fans were used for this study. One active and one inactive lever was presented at the beginning of each session. Above each lever was a stimulus light, and each chamber contained a house light. Infusion pumps (MED Associates) were located outside of each chamber. Experimental events were recorded by MED-PC software on a computer in the experimental room. All operant chambers have been previously described (see Maher et al., 2022; Maher et al., 2021).

#### Experiment 2A

Rats (n=4/sex/condition) were food trained prior to SA procedures (see Maher et al., 2022 for more detail). Rats then underwent fentanyl SA acquisition, whereby 2.5 μg/kg/infusion fentanyl (0.1 mL/infusion), a dose chosen as it is in the middle of the fentanyl dose response curve (Hammerslag et al., 2020; Seaman & Collins, 2021) was delivered across 5.9 s following one response on the active lever (FR-1). Upon an active lever press, lights above both levers were illuminated and a tone (2900 Hz) was presented simultaneously with drug infusion. Infusions were followed by a 20-s dark timeout period, during which active lever responses were recorded but produced no consequences. An inactive lever was present at all times but produced no consequences when pressed. All sessions lasted 2 hr. Daily xylazine injections occurred 15 min prior to SA sessions following the first 10 days of fentanyl SA acquisition, and continued for 8 consecutive days.

#### Experiment 2C

Rats underwent food training followed by fentanyl SA using the same methodology described in Experiment 2A, with or without decreasing doses of xylazine in the fentanyl infusion (3 days each of 0.5, 0.15, and 0.05 mg/kg/infusion xylazine, in descending order with 3 days at each dose).

### PK Methodology (Experiment 2B)

Xylazine HCl was dissolved in saline and given to fasted rats by intravenous bolus at 1.10 mg/kg. Plasma and brain samples were collected at 5 different time points (0.25, 0.5, 1, 2, 4 h). Tissues were homogenized with 9 volumes (w:v) of homogenizing solution (15 mM PBS (pH 7.4): MeOH=2:1). Into each 40 μL aliquot, 400 μL of IS1 (6 in 1 internal standard in ACN including 100 ng/mL of labetalol, tolbutamide, verapamil, dexamethasone, glyburide, and celecoxib) was added, and the resulting mixture was vortex-mixed for 10 min at 800 rpm and centrifuged for 15 min at 3220 × g, 4 °C. Then 50 μL of supernatant was aliquoted and followed by centrifuging for 15 min at 3220 × g, 4 °C before the LC-MS/MS analysis.

### Statistical Analysis

Data for SA infusions and lever press discrimination (active, inactive) across sessions (Experiments 2A, 2C) and withdrawal signs (Experiment 1, 2C) were collected and analyzed using linear mixed effects (LME) modeling using JMP software from SAS. Tukey’s HSD was used as post-hoc pairwise comparisons on nominal variables where appropriate (α = 0.05). For Experiment 2C, data for the intravenous xylazine dose response were averaged across the last 2 sessions at each xylazine dose, converted to percent fentanyl alone as a function of biological sex, and analyzed using LME. Intake from sessions 2, 3, 5, 6, 8, and 9 (the same sessions used for the xylazine dose response in the co-SA groups) was averaged from the fentanyl SA alone male and female groups in Experiment 2C and analyzed via one-tailed t-test to ensure no differences existed between groups prior to utilizing these data as the control for xylazine/fentanyl intake in each sex. Subject was treated as a random factor for all analyses. The total number of infusions per animal during acquisition was recorded per session and compared across groups using LME. Lever discrimination was assessed as the ratio of active/active+inactive lever presses (including infusion-earning and timeout active lever presses, as well as all inactive lever presses), employing our typical criteria for SA acquisition (discrimination above 66.67%) (Maher et al., 2022). PK results were analyzed by a non-compartmental model and Phoenix WinNonlin 6.3 software. A graph was plotted with concentration against time, and data are presented as mean ± standard error of the mean (SEM) from n=2/timepoint. All graphing was performed in Prism 8.1 (GraphPad Software, San Diego, CA). Data from all behavioral experiments are represented as mean ± SEM where appropriate. Transparency and openness guidelines: Materials are available upon request.

## Results

### Experiment 1: Chronic Xylazine Treatment Sex-Specifically Elicits Somatic Signs of Withdrawal

An experimental timeline is displayed in **Figure 1A**_**1**_. We compared somatic signs of withdrawal following chronic experimenter-delivered xylazine or vehicle (saline) injections. LME revealed no significant main effect of treatment group (F_(1,10)_ = 1.93, p > 0.05) or timepoint (F_(1,10)_ = 1.34, p > 0.05), but there was a significant group x timepoint interaction (F_(1,10)_ = 16.46, p < 0.05; **Figure 1B**_**1**_). When broken down by sex, LME analysis revealed no significant main effects of group, sex, or timepoint (p’s > 0.05). However, significant group x timepoint (F_(1,8)_ = 50.10, p < 0.05), sex x timepoint (F_(1,8)_ = 9.45, p < 0.05), and group x sex x timepoint (F_(1,8)_ = 12.99, p < 0.05) interactions were found, indicating that females demonstrated significantly more severe somatic signs of withdrawal relative to males and had a different time course of withdrawal (**Figure 1B**_**2**_).

**Figure 1.**
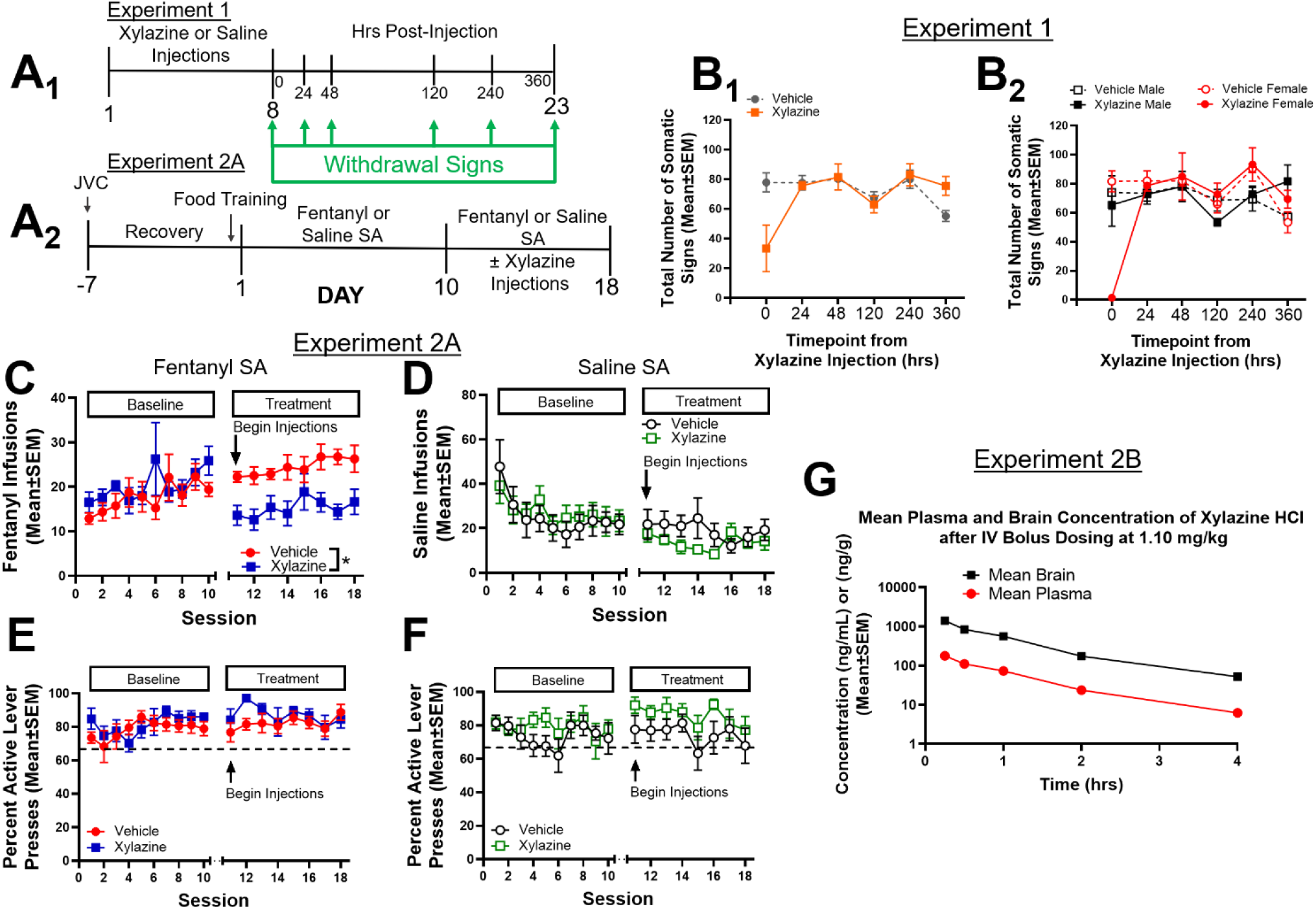
Xylazine suppresses fentanyl but not saline intake during SA. Experimental timelines for Experiment 1 **(A**_**1**_) and Experiment 2A **(A**_**2**_**). (B**_**1**_) Repeated experimenter-delivered xylazine lowers somatic signs of withdrawal acutely, which dissipates within 24 hrs (n=6/group). **(B**_**2**_) Reductions in somatic signs immediately following repeated xylazine injections are driven by females but not males (n=3/sex/group). **(C)** In Experiment 2A, no baseline differences in consumption were found between future xylazine or saline treatment groups. Experimenter-delivered xylazine pretreatments reduced consumption of fentanyl (**C**; \**p*<0.05, the main effect of treatment on sessions 11-18) but not saline during SA (n=8/group) **(D)**. Xylazine pretreatments did not impair lever discrimination during **(E)** fentanyl or **(F)** saline SA, and rats demonstrated percent active lever pressing above 66.67% (dotted lines in E and F). **(G)** Mean brain (ng/g) and plasma (ng/mL) concentration of xylazine after intravenous (IV) bolus dosing in Experiment 2B (n=2/timepoint).

### Experiment 2A: Xylazine Pretreatment Decreases Fentanyl but not Saline SA

Next, we determined the impact of experimenter-delivered xylazine on fentanyl or saline SA and lever discrimination (see experimental timeline in **Figure 1A**_**2**_). LME analyses revealed no differences in baseline fentanyl SA infusions during acquisition (examined across the 10 days of baseline acquisition prior to xylazine or saline pretreatments; **Figure 1C**, left x-axis). Specifically, no main effects of treatment group (F_(1,12)_ = 0.92, p > 0.05) or sex (F_(1,12)_ = 0.06, p > 0.05) were found. There was a significant main effect of the session (F_(1,12)_ = 20.65, p < 0.05), indicating that intake increased across acquisition sessions, but no interactions were found (p’s > 0.05). LME was conducted separately for baseline saline infusions as a function of future treatment groups across sessions. There was only a significant main effect of the session (F_(1,12)_ = 6.12, p < 0.05), indicating that baseline saline intake did not differ between future treatment groups, and saline infusions decreased across sessions (**Figure 1D**, left x-axis). Next, fentanyl infusions during the 8 xylazine or saline treatment sessions (sessions 11-18) were analyzed via LME (**Figure 1C**, right x-axis). A significant main effect of treatment group (F_(1,12)_ = 12.16, p < 0.05) and session (F_(1,11.61)_ = 7.81, p < 0.05), were found, but no effects of sex or group x sex, sex x session, or group x sex x session interactions were found (p’s > 0.05). Finally, saline infusions during the 8 xylazine or saline treatment sessions (11-18) were analyzed via LME (**Figure 1D**, right x-axis). There were no significant main effects or interactions (p’s > 0.05). Together, these results indicate that xylazine pretreatment reduced fentanyl but not saline SA following acquisition, but not in a sex-specific fashion.

Next, lever discrimination during the first 10 sessions (pre-treatment) was analyzed via LME for fentanyl and saline SA separately. For the fentanyl SA groups, LME revealed no significant main effects or interactions (p’s > 0.05; **Figure 1E**, right x-axis). For the saline SA groups, LME also revealed no significant main effects or interactions (p’s > 0.05; **Figure 1F** left x-axis). Lever discrimination was then assessed during the 8 treatment sessions (sessions 11-18) for fentanyl and saline separately. LME revealed no significant main effects or interactions in the saline SA groups (p’s > 0.05; **Figure 1F**; right x-axis). Together, all rats met our SA criterion (e.g., above 66.67% active presses) regardless of treatment, condition, sex, or session, indicating that all rats accurately discriminated the active from the inactive levers during fentanyl and saline SA.

### Experiment 2B: Intravenous Xylazine PK Results

Before introducing xylazine into intravenous fentanyl infusions in Experiment 2C, we first conducted a PK study to determine the half-life (t_1/2_) of intravenous xylazine and the ability of intravenous xylazine to achieve quantifiable levels in brain and plasma. In brain and plasma, t_1/2_ was 0.865 and 0.838 hrs, respectively, following an intravenous bolus high dose (1.10 mg/kg) of xylazine. The data in **Figure 1G** show that xylazine was present in the brain and plasma following bolus high-dose intravenous dosing. Clinical signs from the high dose included an unsteady gate and decreased activity, which resolved within 1 hr post-intravenous administration.

### Experiment 2C: Intravenous Xylazine Reduces Fentanyl SA and Impacts Fentanyl Withdrawal Symptomatology

Rats next underwent intravenous xylazine/fentanyl SA based on the PK results from Experiment 1C (see timeline in **Figure 2A**). Infusions as a function of xylazine dose are shown in **Figure 2B**. LME analyses were conducted using percent fentanyl SA alone and revealed a significant main effect of xylazine dose (F(_1,22_) = 75.37; p < 0.05; **Figure 2C**), and a main effect of sex (F(_1,22_) = 6.53; p < 0.05), but no sex x dose interaction (p > 0.05). Next, LME was conducted on percent active lever presses from both fentanyl SA alone and xylazine/fentanyl co-SA, and no main effects of sex, session, xylazine dose, or interactions were found (p’s > 0.05; **Figure 2D**). All groups showed lever discrimination above 66.67%, demonstrating that all rats learned the task regardless of the xylazine dose.

**Figure 2.**
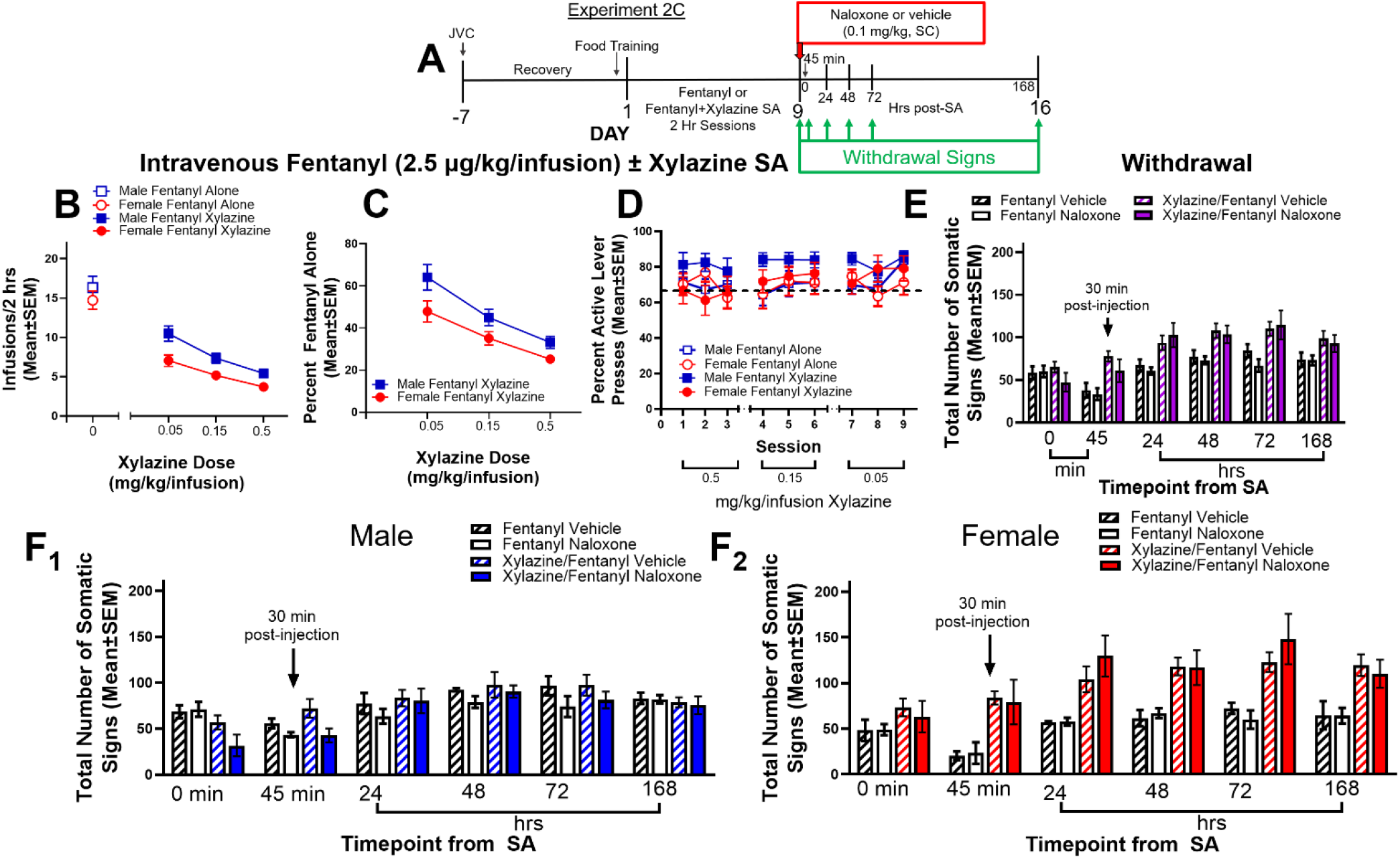
Intravenous xylazine/fentanyl co-SA reduces the intake in a dose-dependent fashion and induces a sex-specific withdrawal syndrome that is resistant to acute precipitation of withdrawal by naloxone. **(A)** Experimental timeline for Experiment 2C. **(B)** Raw infusions in male and female rats from either fentanyl SA without xylazine in the infusion or with varying doses of xylazine in the fentanyl infusion (n=6/sex/group). **(C)** When evaluated compared to fentanyl SA alone, intravenous xylazine dose-dependently reduced intake during xylazine/fentanyl SA (n=6/sex/group). **(D)** Intravenous xylazine/fentanyl SA did not impair lever discrimination at any xylazine dose tested (dotted line at 66.67%; n=6/sex/group). **(E)** Somatic signs of withdrawal did not differ between naloxone and vehicle-treated groups following xylazine/fentanyl co-SA (fentanyl alone: n=6/group; xylazine/fentanyl co-SA: n=12/group). When broken down by sex **(F**_**1**_; n=6/sex/group), naloxone did not precipitate signs of withdrawal in males regardless of whether xylazine was included in the fentanyl infusion, and there were no differences in somatic signs of withdrawal in males regardless of whether fentanyl was self-administered alone or with xylazine in the infusion; however, **(F**_**2**_) xylazine/fentanyl co-SA females demonstrated higher somatic signs of withdrawal across timepoints compared fentanyl SA alone, demonstrating that xylazine/fentanyl co-SA elevates withdrawal symptomatology across timepoints in females (\**p*<0.05 indicating significance between fentanyl SA alone and xylazine/fentanyl co-SA female groups; n=6/sex/group).

Next, we conducted LME analyses on somatic signs of withdrawal from both fentanyl and xylazine/fentanyl groups following naloxone or vehicle treatment. First, LME was conducted to determine if naloxone precipitated withdrawal following fentanyl SA alone or xylazine/fentanyl co-SA. LME analysis indicated a significant main effect of timepoint (F(_1,32.01_) = 66.71; p < 0.05) and group (F(_1,31.97_) = 7.6; p < 0.05), but no group x timepoint interaction (p > 0.05; **Figure 2E**). Because there is one clinical study showing that women display more severe opioid withdrawal compared to men (Ware et al., 2022), we hypothesized that females would display more severe withdrawal symptomatology compared to males when analyzed by sex. LME of fentanyl alone and xylazine/fentanyl co-SA did confirm significant main effects of group (F_(1,27.76)_ = 10.08; p < 0.05) and timepoint (F_(1,28)_ = 64.57, p < 0.05), as well as a significant group x sex interaction (F_(1,27.76)_ = 10.01; p < 0.05; see data broken down by sex in **Figure 2F**_**1**_ (males) and **Figure 2F**_**2**_ (females)). Post-hoc analyses confirmed that withdrawal signs from fentanyl SA alone and xylazine/fentanyl co-SA were significantly different only in females. Together, these results suggest that xylazine/fentanyl co-exposure may uniquely alter withdrawal symptomatology across time in females relative to males, but that this is not altered by acute naloxone treatment.

## Discussion

We are the first to show that intravenous xylazine adulteration of fentanyl uniquely alters fentanyl SA and withdrawal symptomatology in a rodent model of xylazine/fentanyl co-SA. We further demonstrate that repeated xylazine induces its own unique withdrawal syndrome, but only in female rodents whereby there is an acute suppression of somatic signs of withdrawal. Here, we establish a rat model of intravenous xylazine adulteration of fentanyl SA. We show that both experimenter-delivered or self-administered xylazine reduced fentanyl intake. When self-administered, intravenous xylazine decreased fentanyl SA, and xylazine dose-dependently decreased fentanyl intake in both sexes, though examination of additional doses of xylazine is warranted. In general, examination of naloxone’s effects 30 minutes after administration did not reveal a large increase in somatic signs of withdrawal (at the 45 min post-SA timepoint), regardless of whether xylazine was included in the fentanyl infusion. This could be due to the potential need for higher naloxone doses or repeated dosing of naloxone to fully reverse fentanyl-related effects such as respiratory depression (given fentanyl’s higher potency for the mu opioid receptor as compared to other opioids [Fairbairn et al., 2017; Hedrick et al., 2021]); however, additional naloxone doses and/or repeated naloxone dosing are needed to fully evaluate these possibilities. Surprisingly, animals in the xylazine/fentanyl co-SA groups demonstrated higher somatic signs of withdrawal (regardless of naloxone treatment) as compared to animals self-administering fentanyl alone (**Figure 2E**), and this appeared to be driven by females (as shown in **Figure 2F**). Together, these results indicate that regardless of sex or xylazine adulteration of fentanyl, naloxone was unable to acutely increase somatic signs of withdrawal but that xylazine/fentanyl co-SA may induce a unique and elevated withdrawal syndrome in females.

In addition to SA of the drug combination and evaluation of withdrawal, this study evaluated outcomes as a function of biological sex. There is a dearth of information at the clinical level of analysis regarding sex differences in aspects of the opioid use disorder (Huhn et al., 2018), and no information on sex-specific impacts of xylazine/fentanyl co-SA. Consistent with one recent clinical study, which demonstrated that women report more severe opioid withdrawal than men (Ware et al., 2022), we found that somatic signs of withdrawal were more pronounced in females as compared to males following xylazine/fentanyl co-SA. Further, females may have been more sensitive to the motor suppressive effects of xylazine when administered repeatedly as compared to males, however, additional studies are needed to fully evaluate possible locomotor-suppressing effects of xylazine in females. Withdrawal was not different between experimenter-administered xylazine and vehicle-treated males, indicating that xylazine itself (at the dose examined) did not induce a withdrawal syndrome in males. Although additional research is needed, our results also show that naloxone was ineffective at significantly increasing signs of withdrawal 30 min post-treatment regardless of sex. Given that the half-life of naloxone in rats is ∼30 min (Ngai et al., 1976), we postulate that the sex-specific protracted withdrawal effects seen here following xylazine/fentanyl co-SA were not due to naloxone treatment.

Given the sex-specific withdrawal effects following xylazine/fentanyl co-SA, it is possible that xylazine may impact the ability of fentanyl to undergo adipose-related excretion patterns. Fentanyl is highly lipophilic molecule (Roy & Flynn, 1988) that is redistributed to adipocytes, where it is subject to an extended “secondary peaking” excretion pattern (Comer & Cahill, 2019; Huhn et al., 2020). Our data show that only xylazine/fentanyl co-SA females demonstrated elevated withdrawal signs across timepoints (regardless of naloxone administration), raising the possibility that the addition of xylazine to fentanyl infusions may alter secretion of fentanyl from adipose tissue in a sex-specific fashion. Although not studied here, prior literature indicates that clonidine, which is an agonist of alpha 2a (α2a) adrenergic receptors similar to xylazine, inhibits brown adipose tissue (BAT) sympathetic nerve activity and thermogenesis (Madden et al., 2013). Thus, it is possible that xylazine impacted adipose functions via agonism of α2a receptors. It is further possible that xylazine and fentanyl were sequestered in BAT, and subsequently released across withdrawal days (deleterious effects on glucose homeostasis have been found after exposure to lipophilic organic pollutants during weight loss [Baker et al., 2013]). Given that females likely had more BAT than males when housed at room temperature, it is possible that withdrawal from the xylazine/fentanyl combination resulted in greater loss of BAT in females than males, which may have allowed these drugs to be released from adipose stores and have impacts on withdrawal. However, future studies are needed to confirm these possibilities.

The present studies are limited in that they did not include a saline/xylazine SA group, did not manipulate fentanyl dose, did not evaluate withdrawal in Experiment 2A, and did not provide opportunity for rats to acquire fentanyl SA prior to intravenous xylazine administration. The dose of fentanyl utilized in the current studies was selected because it is in the middle of the dose-response curve (Hammerslag et al., 2020; Seaman & Collins, 2021). However, without additional doses of fentanyl, it is not possible to determine if xylazine induces a leftward, rightward, or downward shift in the fentanyl dose-response curve. An additional limitation is that only three doses of xylazine were tested in fentanyl SA, and thus additional lower doses should be examined to further evaluate the xylazine/fentanyl dose-response curve. We also only utilized short access sessions, thus it is possible that the lower consumption of fentanyl due to the inclusion of xylazine may not have resulted in a robust withdrawal syndrome, and this may underlie the lack of precipitated withdrawal following naloxone treatment. Finally, one other limitation to these studies is the inclusion of a saline SA group rather than a food control group, given the low rates of responding due to saline self-administration and the possibility of a floor effect with xylazine administration in Experiment 2A. The reason for this possible limitation is that it may be important to match rates of behavior between fentanyl and control groups prior to xylazine treatment, given rate-dependency in which responding following a treatment is dependent upon the baseline rate of behavior (Dews, 1958; Wenger & Dews, 1976). Regardless of these limitations, the present study provides critical and entirely novel information regarding SA of and withdrawal from an emerging problematic pattern of polysubstance use and provides a foundation upon which future studies can refine these methodologies.

### Translational Implications and Future Directions

The present studies demonstrate novel findings regarding the impacts of xylazine on fentanyl-induced behaviors, including SA and withdrawal, and provide a foundation for future studies to expand upon in this area. For example, one future evaluation should examine these outcomes as a function of the estrous cycle phase, especially given the unique behavioral patterns that emerged here in females. Future studies could also address whether there are specific neurobiological adaptations following xylazine/fentanyl co-SA, which may underlie the current behavioral findings.

Clinical studies on xylazine/fentanyl co-SA provide epidemiological evidence of this novel and problematic pattern of co-SA (e.g., Bowles et al., 2021; Friedman et al., 2022). However, no studies to date have evaluated the behavioral or neurobiological impacts of repeated xylazine/fentanyl co-SA, and none have empirically demonstrated how co-SA of these substances changes fentanyl consumption and/or induces a unique sex-specific withdrawal syndrome. Thus, the present study represents the first empirical evidence demonstrating these behavioral effects and lays a foundation upon which preclinical non-human and human clinical studies can further evaluate this pattern of polysubstance use.

